# Disrupted calcium dynamics in reactive astrocytes occur with endfeet-arteriole decoupling in an amyloid mouse model of Alzheimer’s disease

**DOI:** 10.1101/2025.01.24.634584

**Authors:** Blaine E. Weiss, John C. Gant, Ruei-Lung Lin, Jenna L. Gollihue, Susan D. Kraner, Edmund B. Rucker, Yuriko Katsumata, Yang Jiang, Peter T. Nelson, Donna M. Wilcock, Pradoldej Sompol, Olivier Thibault, Christopher M. Norris

## Abstract

While cerebrovascular dysfunction and reactive astrocytosis are extensively characterized hallmarks of Alzheimer’s disease (AD) and related dementias, the dynamic relationship between reactive astrocytes and cerebral vessels remains poorly understood. Here, we used jGCaMP8f and two photon microscopy to investigate Ca^2+^ signaling in multiple astrocyte subcompartments, concurrent with changes in cerebral arteriole activity, in fully awake eight-month-old male and female 5xFAD mice, a model for AD-like pathology, and wild-type (WT) littermates. In the absence of movement, spontaneous Ca^2+^ transients in barrel cortex occurred more frequently in astrocyte somata, processes, and perivascular regions of 5xFAD mice. However, evoked arteriole dilations (in response to air puff stimulation of contralateral whiskers) and concurrent Ca^2+^ transients across astrocyte compartments were reduced in 5xFAD mice relative to WTs. Synchronous activity within multi-cell astrocyte networks was also impaired in the 5xFAD group. Using a custom application to assess functional coupling between astrocyte endfeet and immediately adjacent arteriole segments, we detected deficits in Ca^2+^ response probability in 5xFAD mice. Moreover, endfeet Ca^2+^ transients following arteriole dilations exhibited a slower onset, reduced amplitude, and lacked relative proportionality to vasomotive activity compared to WTs. The results reveal nuanced alterations in 5xFAD reactive astrocytes highlighted by impaired signaling fidelity between astrocyte endfeet and cerebral arterioles. The results have important implications for the mechanistic underpinnings of brain hypometabolism and the disruption of neurophysiological communication found in AD and other neurodegenerative conditions.

**Significance Statement:** Astrocytes are an essential component of the neurovascular unit. Chronically reactive astrocyte phenotypes are mechanistically linked to deleterious features of Alzheimer’s disease (AD) including impaired cerebral blood flow, hypometabolism, and synapse dysfunction/loss. Here, we show that reactive astrocytes in a fully awake mouse model of AD-like amyloid pathology are spontaneously hyperactive, exhibit impaired functional connectivity, and respond to dilations in immediately adjacent arterioles with poor fidelity. The results reveal a key point of communication breakdown between the brain and the cerebrovasculature.

## INTRODUCTION

The neurovascular unit, comprised of cerebral blood vessels and multiple neural cell types is essential for promoting brain health(Iadecola, 2017; Schaeffer and Iadecola, 2021). Chronic neurodegenerative diseases like Alzheimer’s disease (AD) are characterized by pathognomonic proteinaceous deposits that occur alongside cerebrovascular dysfunction and hypometabolism(Veitch et al., 2019). Most of the cerebral vasculature is ensheathed by astrocyte endfeet specializations, while other astrocyte processes are in direct contact with many, if not most, excitatory synapses(Verkhratsky and Nedergaard, 2014; Pivoriunas and Verkhratsky, 2021). Astrocytes also contain the requisite machinery for taking up, processing, and releasing metabolites, and therefore represent a critical cellular liaison between blood and brain. In the course of AD and related dementia disorders (ADRDs), astrocytes become reactive and exhibit complex loss- and gain-of-function phenotypes(Escartin et al., 2021; Price et al., 2021; Liddelow et al., 2024). These clues have led us and others to hypothesize that astrocytes play key roles in the deleterious cerebrovascular phenotypes of chronic neurodegenerative conditions(Beard et al., 2021; Price et al., 2021; Stackhouse and Mishra, 2021; Takahashi, 2022).

Recent studies have investigated Ca^2+^ signaling as a proxy for the altered function of reactive astrocytes in AD/ADRD mouse models(Takano et al., 2007; Kuchibhotla et al., 2009; Delekate et al., 2014; Lines et al., 2022; Shah et al., 2022; Kelly et al., 2023; Lee et al., 2023; Sompol et al., 2023). Findings have been mixed with some reports showing augmented Ca^2+^ signaling in AD mouse models, characterized by elevations in spontaneous Ca^2+^ transient frequencies and amplitudes, while others have shown no change or reductions in these parameters. To date, only a few reports on intact AD/ADRD mouse models have described Ca^2+^ signaling in the endfeet(Delekate et al., 2014; Kelly et al., 2023; Lee et al., 2023), which are the most proximal astrocyte subcompartments relative to the cerebral blood vessels. In these studies, endfeet Ca^2+^ was either assessed in the absence of concurrent vascular measures(Kelly et al., 2023), or during the administration of general anesthesia(Delekate et al., 2014; Lee et al., 2023), which can greatly suppress astrocyte signaling and/or alter vascular tone during brain stimulation(Thrane et al., 2012; Gao et al., 2017).

Here, we used the genetically-encoded Ca^2+^ indicator jGCaMP8f and intravital two photon microscopy to investigate astrocyte Ca^2+^ transients in multiple cellular compartments of fully awake 5xFAD mice (characterized by robust parenchymal brain amyloid pathology) and age-matched wild type littermates. Astrocyte Ca^2+^ signaling in individual subcompartments and astrocyte networks, concurrent with dynamic responses in penetrating arterioles, was assessed in the barrel cortex at rest and during air puff whisker stimulation of the contralateral vibrissae. Using a novel custom vascular modeling application, Localized Analysis of Vascular Astrocytes (LAVA), we also directly assessed temporal and proportional relationships between astrocyte endfeet Ca^2+^ transients and immediately adjacent arteriole segments. Our findings describe novel (dys)functional phenotypes of reactive astrocytes and may offer new insights into how energy coupling breaks down during AD and related dementias.

## MATERIALS AND METHODS

### Mice

Male and female 5XFAD and wild type (B6-SJL)(Oakley et al., 2006) mice were purchased from The Jackson Laboratory (Cat #) and bred to produce hemizygous and littermate control F2 mice. Mice were housed in standard laboratory cages (IVC) under 12 h light/dark cycles in a pathogen-free environment in accordance with University of Kentucky guidelines. Mice had access to food and water *ad libitum*. All animal procedures were conducted in accordance with the National Institutes of Health Guide for the Care and Use of Laboratory Animals and were approved by University of Kentucky Institutional Animal Care and Use Committees.

### Adeno-associated virus (AAV) reagents

cDNAs for jGCaMP8f(Tran et al., 2023) were encoded in vectors downstream of the human GFAP promoter (GFA104 ABCD9). Plasmids were sent to Charles River Laboratories for packaging into AAV2/5 capsids. When injected into adult mice, AAV-GFA104 vectors drive expression of target transgenes selectively in astrocytes. Mice were injected with AAV2/5-GFA104-JCaMP8f into barrel cortex (see below) at approximately six months of age to detect and quantify astrocytic Ca^2+^ transients.

### AAV injection and cranial window implantation

Surgical techniques were performed as described in our previous studies(Case et al., 2023; Sompol et al., 2023) with some modifications. Briefly, mice were anaesthetized by 3% isoflurane (SomnoSuite, Kent Scientific) in an induction box and head-fixed to a stereotaxic frame. Anesthesia was maintained by continuous isoflurane inhalation (1.5-3%, via nose cone) during surgery. The injection was marked over the barrel cortex at 1.5mm AP, ± 3mm ML. The injection site was used as the center point of a 3mm diameter craniotomy for cranial window implantation. The freed circular skull fragment was carefully removed with tweezers. AAV (10^13^ IFU/mL) was delivered at a rate of 0.2 µL/min (total 2uL) into the brain using a Hamilton syringe and microinjector (Stoelting). After injection, needle remained in place for an additional 2 min prior to retraction. Then a glass cranial window, made from 3 and 4 mm glass coverslips combined with optical glue, was carefully placed over the craniotomy with the smaller coverslip down, to cap the exposed brain in the skull opening. The window was stabilized, and affixed to the skull along with a stainless-steel head mount (Protoscience Solutions, Lexington, KY) using dental acrylic. Analgesic was injected subcutaneously, and animals recovered from anesthesia before return to their home cages.

### Two photon microscopy

Mice were anesthetized and affixed to the imaging platform via a custom made fixture (Protoscience Solutions, Lexington, KY) on a Scientifica Hyperscope utilizing a tunable InSight X3 laser (Spectra-Physics), ranging from 680 to 1300nm, and a 16X, 0.8 NA, 3 mm working distance objective lens (Nikon Instruments), and GaAsP photomultipliers (Hamamatsu Photonics). ScanImage acquisition software (MBF Biosciences) was used to drive microscope components and data acquisition from within MATLAB (MathWorks). Mice were aligned under the objective lens and received single retroorbital injections of rhodamine dextran dye (500 kDa, 5% w/v in saline) to visualize cerebral vasculature. A small air pump was gated by a solenoid control valve and connected to the vDAQ controlled through ScanImage. (MBF Biosciences). Anesthesia was removed, and imaging commenced once mice were fully alert and excitable by air puff stimulation (20 min or less).

### Intravital Imaging of Vascular Dynamics and Astrocyte Ca^2+^ Signaling

Imaging sessions began by surveying the cranial surface for expression of GCaMP8f indicator. Once the field of view was selected, a baseline 5-minute recording of vascular and astrocyte activity was acquired followed by a series of timed whisker stimulation trials. Stimulation consisted of 10 second air puff trains (9 Hz) to the contralateral vibrissae, controlled by a custom software trigger integrated into the ScanImage software’s frame clock. Videos were captured that contained several astrocytes, cerebral capillaries, and arterioles within the field of view. A minimum three minute interval separated stimulation trials. At the conclusion of the experiment, mice were then lightly anesthetized for removal from the imaging platform, and placed back into their home cages.

### Image Analysis

Experimenters were blinded from genotype during all analyses, which were conducted using in-house Matlab algorithms described here. First, files containing time lapse videos (*i.e.* .tifs) were motion corrected by the widely used NoRMCoRRE algorithm(Pnevmatikakis and Giovannucci, 2017). Metadata containing scan speed, spatial dimensions, and stimulation timing were extracted for common use by analyses applications. Event based detection methods were used to identify astrocytes in the field of view (FOV). Further segmentation of astrocyte compartments such as soma and endfeet was accomplished through convolutional filtering and masking techniques that accentuate astrocyte features. For astrocyte endfeet, the perivascular space was masked prior to event based detection. In depth analyses of vascular motion and spatially paired astrocyte calcium signaling were characterized using our custom MATLAB-based application called LAVA. Vascular motion was assessed by linearizing vessels of interest to better define spatial alignment between arteriole segments and endfeet compartment activity. Cross sectional vectors placed over arterioles of interest were used to determine cross sectional area and to sample fluorescence values of adjacent endfeet. The resulting “dataset” provides spatial-temporal information that can be used to quantify dynamic relationships between vascular tone, motion, and/or Ca^2+^ fluorescence, which, in turn, can offer insights into the directionality of vascular tone fluctuations and potential Ca^2+^ mediated changes. LAVA can also be used to correlate this activity in relation to other ROI segmentations (*e.g.* somata and capillary endfeet signals mentioned previously and not shown in this report), for predefined trial phases (*i.e.* before, during, and post-stimulation). The workflow for the application is as follows: In each FOV, dilating and contracting vessels-of-interest are identified and vessel boundaries are outlined. Vessels are then binarized and archived. The resulting data structure contains a total volume time series vector, and fully cross-sectioned and linearized vascular and Ca^2+^ channel image stacks. User defined perivascular ROIs are then identified from the linearized and scaled pseudo images of fluorescence changes along the vessel-of-interest. Results from vascular motion measurements are optionally used to update ROI positions for arteriole endfeet where vascular movement contaminates extracted Ca^2+^ signals. The application then extracts and saves the resulting time series vectors from these ROIs, and calculated maximum relative fluorescence changes. Correlations between endfeet Ca^2+^ signals and vascular activity are quantified by Pearson’s r. Once ROI-based endfoot analyses are completed, LAVA conducts within vessel comparisons of fluorescence time series data and local vascular tone, by correlations within cross-sectional vectors. Video outputs of processed data are available from within LAVA and are demonstrated in 3.

### Statistics

Most Ca^2+^ transient and vessel property parameters were compared across groups using unpaired student t-tests. Arteriole diameter changes before and during air puff stimulation were evaluated with repeated measure ANOVA, followed by simple effects tests. Correlations between time series data were determined by Pearson’s r tests. Specifically, for network synchronicity analysis, events were cropped to only the rise phase of Ca^2+^ event transients before statistical analysis. Roughly equal numbers of males and females were used per genotype condition (WT: 5M & 6F, 5xFAD: 5M 9F). The total group numbers for animals were 11 WT and 14 5xFAD. At least two FOVs per mouse were used, and three replicate trials each for stimulation experiments. Not all parameters could be cleanly measured fro13m each mouse (*e.g.* interruption in imaging session, lack of identifiable arterioles, poor expression of Ca^2+^ indicator), resulting in exclusion from statistical analyses. Exclusion was performed before decoding of transgene groups. Unless, otherwise noted, statistical samples shown are average values obtained within each mouse. Significance for all statistical tests was set at *p* < 0.05. For most parameters evaluated, there were no significant sex differences in either transgene group and so males and females were grouped together.

### Software Accessibility

The software described herein and related documentation is available for download via its Github repository. <u>Localized Analysis of Vascular Astrocytes (LAVA)</u> © 2024 by <u>Blaine Weiss</u> is licensed under <u>CC BY-NC-SA 4.0</u> and the Norris Laboratory webpage https://norris.createuky.net/.

## RESULTS

### Spontaneous Ca^2+^ transients in 5xFAD mice are smaller, but occur more frequently in astrocyte somata and processes

Two photon imaging was used to assess astrocyte Ca^2+^ transients in barrel cortex of fully awake mice at a depth of 150-300 µm. Spontaneous Ca^2+^ transients were first assessed in the absence of movement across a 5 min time period (see Extended data 1), in which no external stimulation was administered. Regions of interest (ROIs) representing astrocyte somata and processes were segmented across the field of view (FOV) (Figure 1A-D) and a convolution-based event detection technique was used to detect significant events over baseline Ca^2+^ levels. Ca^2+^ transients (ΔF/F) in representative ROIs from a single FOV in a WT and 5xFAD mouse are shown in Figure 1E. Ca^2+^ transients in both cellular compartments occurred more frequently in 5xFAD mice, especially in astrocyte processes (Figure 1F somata, p = 0.06; Figure 1K processes, p = 0.02). However, transient amplitudes in 5xFAD astrocytes were also smaller (Figure 1G somata, p = 0.03; Figure 1L processes, p = 0.03) with trending reductions in area under the curve (AUC) measures (Figure 1I somata, p = 0.05; Figure 1N processes, p = 0.07). No differences in rise time properties were observed (Figures 1H and 1M), but transient decay was accelerated in the 5xFAD group (Figure 1J somata, p = 0.02; Figure 1O processes, p = 0.03).

**Figure 1.**
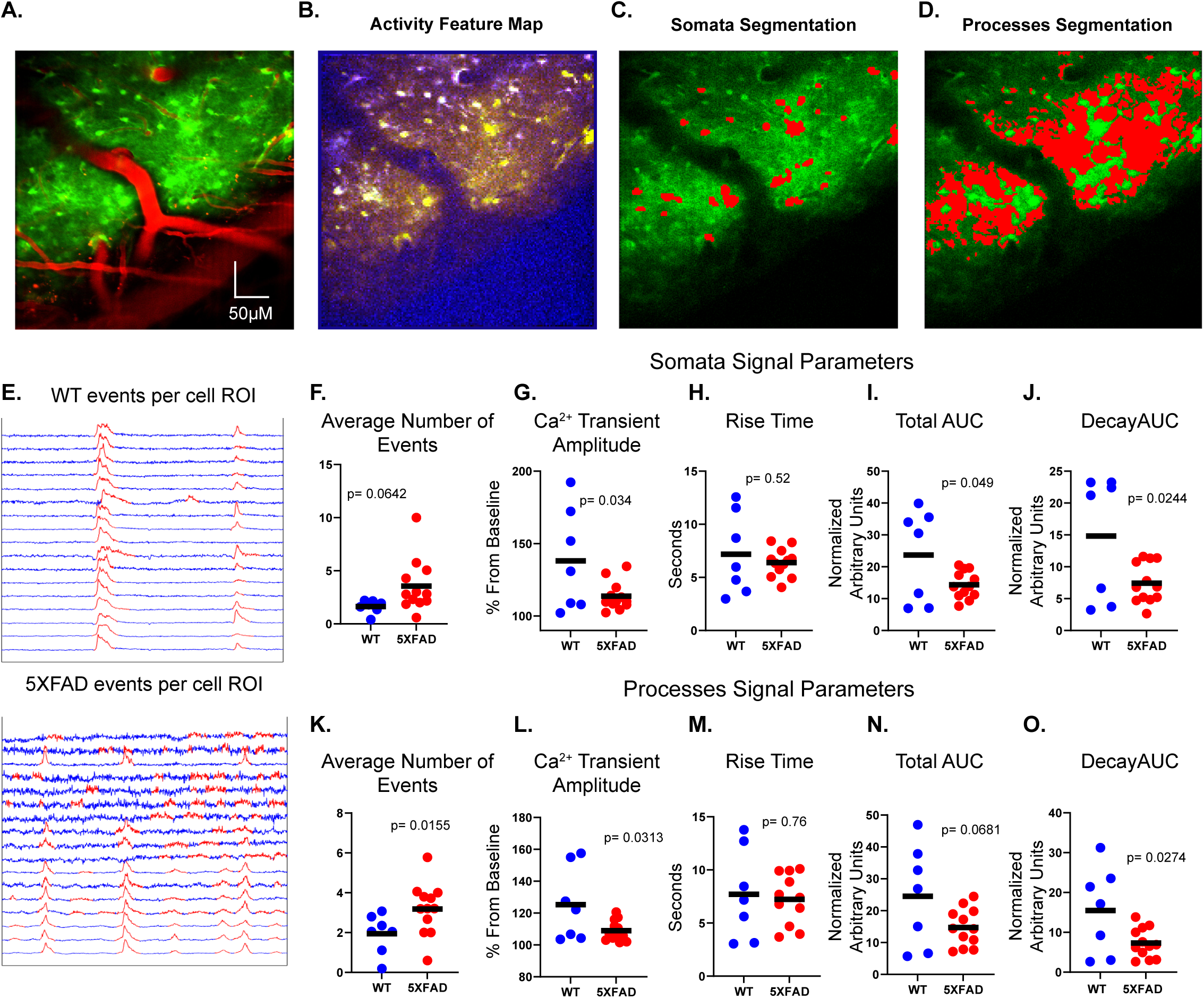
Segmentation of astrocyte sub compartments and spontaneous signal analyses in barrel cortex of awake mice. **A**, Two photon micrograph of a cerebral penetrating arteriole and surrounding capillary bed (red) and summated activity map of astrocyte Ca^2+^ signals (green). **B**, Activity feature map of signal properties (amplitude-red, variability-green, and shape-blue) represent the strength of each feature. Signal features for each pixel were inputted into an unsupervised learning algorithm and clustered to identify pixels composing active astrocyte compartments. **C-D**, Somata (C) and fine process (D) segmentation masks used for ROI signal extraction. **E,** Representative spontaneous Ca^2+^ traces extracted from individual soma compartments in FOVs from a WT and a 5xFAD mouse. Regions in red denote where individual Ca^2+^ transients were detected and measured. **F-O**, Scatterplots show Ca^2+^ transient parameters extracted from astrocyte somata (**F-J**) and processes (**K-O**) and averaged across FOVs for WT and 5xFAD mice. Parameters included: average number of Ca^2+^ transients per ROI (**F,K**), Ca^2+^ transient amplitude (**G,L**), rise time (**H,M**), total AOC (**I,N**) and decay (**J,O**). Each plot symbol represents an individual mouse. *p* values derived from two-tailed T tests.

### Evoked dilations in penetrating arterioles are reduced in 5xFAD mice

Functional hyperemia, as assessed by stimulus-evoked cerebral vessel dilations or increased blood flow is impaired in multiple AD/ADRD rodent models including 5xFAD mice, when assessed under general anesthesia(Mughal et al., 2021). To determine if similar deficits occur in fully awake 5xFAD mice, we delivered air puffs (9 Hz, 10 sec) to contralateral vibrissae and measured hyperemic responses in penetrating arterioles of the barrel cortex (Figure 2A-D). Vessels of interest were identified and cross-sectioned to generate a linearized vessel model (Figure 2E, and see Figure 5A-C). Representative two photon images show penetrating arterioles before (Figure 2B) and during (Figure 2C, D) the air puff, while representative time plots illustrate differences in arteriole dilation across WT and 5xFAD mice (Figure 2F). Before-and-after (air puff) scatterplots are shown in Figure 2G and differences determined by repeated measures 2-way ANOVA. No genotype differences were observed for resting (prestimulation) arteriole tone (Figure 2G, round plot symbols). But, while air puff stimulation led to increased arteriole cross sectional area in all animals [Baseline *vs.* Stimulation F(1,24) = 69.3, p <0.001], this increase was significantly smaller in 5xFAD mice (Interaction: F(1,24) = 5.097, p = 0.033, & Figure 2H, p = 0.002) , indicative of a neurovascular coupling deficit.

**Figure 2.**
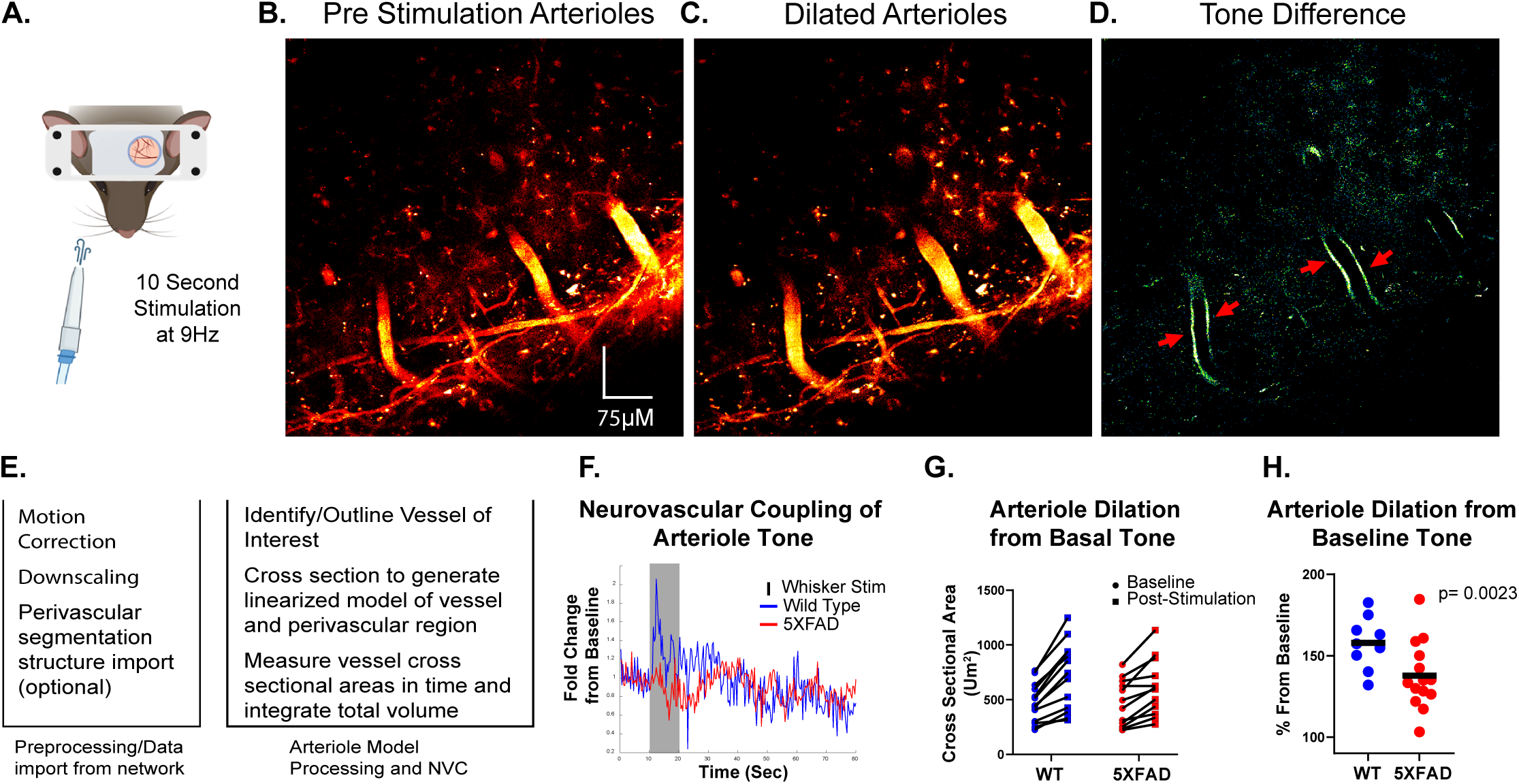
Air puff-evoked arteriole dilations in barrel cortex of awake mice. **A**, Mice received a 10 sec duration 9 Hz air puff train to the contralateral vibrissae. **B-C**, Pseudo-colored two photon micrographs showing penetrating arterioles, before (**B**) and during (**C**) air puff stimulation. **D**, Subtracted image of panels **B** and **C** showing the extent of vessel dilation (red arrows, green vessel segments) during air puff stimulation. **E**, Arteriole modeling and analysis protocol using LAVA application. **F**, Time plots showing representative arteriole diameter changes for WT and 5xFAD mice before, during, and after air puff stimulation (shaded rectangle). **G**, Raw arteriole cross sectional areas measured within each mouse at rest (circles) and during (squares) air puff stimulation. Basal arteriole diameter was comparable across genotypes, and both groups showed significant dilatory responses during air puff delivery [Baseline cross sectional area vs stimulation cross sectional area F(1,24) = 69.3, p <0.001], though dilations were greater in the WT group [Interaction: F(1,24) = 5.097, p = 0.033]. **H**, Scatter plot shows percent arteriole dilation during air puff stimulation for both transgene groups. Each plot symbol represents an individual mouse. *p* values derived from two-tailed T tests.

### Evoked Ca^2+^ activity in astrocytes is impaired in individual cell compartments and across multicell networks in 5xFAD mice

Air puff stimulation led to a large population Ca^2+^ response across astrocytes in the FOV, with highly correlated activity observed in both WT and 5xFAD mice (see Extended data 2 and Figure 3). Figure 3A shows representative extracted Ca^2+^ traces from each ROI in a FOV during a whisker stimulation trial. Ca^2+^ transient kinetics were very similar and synchronous across some ROIs (indicated by asterisks in linearized traces, Figure 3B), while Ca^2+^ transients in other ROIs showed dissimilar kinetics (indicated by pound symbols in linearized traces, Figure 3B). Figure 3C shows representative astrocyte network responses in a WT and 5xFAD mouse with overlaid correlograms. Lines connect co-active ROIs. Line color indicates the degree of synchronicity between each ROI pair (red -yellow is high; blue is low, Pearson’s *r*). Assessment of population responses in astrocyte networks showed reduced synchronicity in the 5xFAD group, especially in astrocyte processes (Figure 3D, somata p = 0.01; Figure 3I, processes p = 0.007 n = 9,13). In addition to network-wide Ca^2+^ responses, we also assessed evoked Ca^2+^ transient parameters from individual cell compartments (somata, Figure 3E-3H; processes, Figure 3J-3M). Individual Ca^2+^ transients were reduced in amplitude in 5xFAD mice (somata, *p* = 0.002, Figure 3E; processes, p = 0.01, Figure 3J), along with trends for reduced AUC measures (somata, p = 0.11, Figure 3G; processes p = 0.08, Figure 3L). The decay time for evoked Ca^2+^ responses was accelerated in the 5xFAD group, especially in astrocyte processes (somata, p = 0.08, Figure 3H; processes, p = 0.03, Figure 3M). No changes in rise time for evoked transients were observed (Figure 3F, 3K). The results show that evoked astrocyte activity, as assessed in individual subcompartments and across astrocyte networks, is impaired in 5xFAD mice.

**Figure 3.**
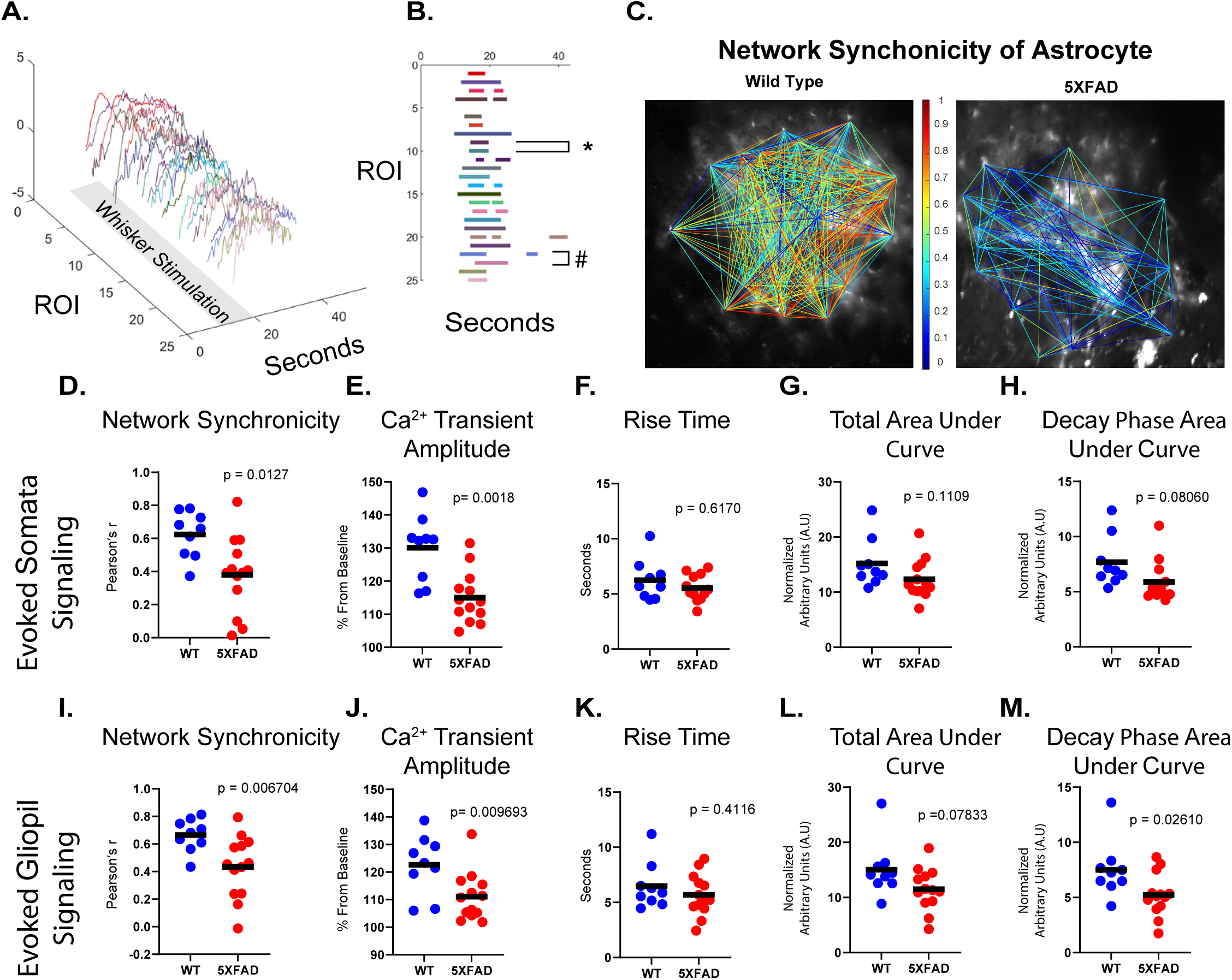
Astrocyte network activation and evoked signal analyses in awake mice. **A**, Individual Ca^2+^ events (shown in different colors) were extracted from unique ROIs in the FOV after whisker stimulation (gray rectangle). **B**, Temporal alignment of Ca^2+^ signaling events from unique ROIs. Each row represents Ca^2+^ events from a unique ROI (y axis). The **\*** symbol exemplifies well synchronized events across ROIs, while the **#** symbol denotes poorly synchronized Ca^2+^ events across ROIs. **C,** Comparison of representative network diagrams of field astrocyte signal synchronization in a WT and 5xFAD mouse (also see Extended data 2). Line colors represent Pearson’s r correlation values. (Negative correlations not shown.) **D-M**, Scatterplots showing evoked astrocyte Ca^2+^ signaling parameters in WT and 5xFAD mice. Each plot symbol represents an individual mouse. Network synchronicity (**D**, **I**) reflects the average Pearson correlation coefficients for all correlated ROI pairs (astrocyte somata, **D**; astrocyte processes, **I**) within the FOV. Ca^2+^ transient amplitude (**E**, **J**), rise time (**F**, **K**), AOC (**G**, **L**), and decay (**H**, **I**) were assessed in individual astrocyte compartments (somata **D-H**; processes **I-M**) and averaged within the in the FOV for each mouse. *p* values derived from two-tailed T tests.

### Changes in perivascular Ca^2+^ signaling in 5xFAD mice

We next assessed perivascular astrocyte Ca^2+^ signaling dynamics, specifically. This entailed activity in the immediate vicinity of all cerebral vessels (arterioles, venules, capillaries) and was enriched for astrocyte endfeet and distal astrocyte process compartments (Figure 4). Representative masking and segmentation steps are shown in Figure 4A and B. Representative activity from multiple perivascular ROIs in the FOV is shown in Figure 4C. In the absence of air puff, spontaneous events occurred in 5xFADs at a higher rate, relative to WTs (Figure 4D, p = 0.02), but tended to be smaller in amplitude (Figure 4E, p = 0.1) and integrated area (Figure 4G p = 0.05), with accelerated decay (Figure 4H, p = 0.02). During air puff stimulation trials, population responses in perivascular regions showed reduced synchronicity in 5xFAD mice (Figure 4I, p = 0.004). Evoked Ca^2+^ transients in perivascular spaces were also smaller in amplitude (Figure 4J, p = 0.001) and integrated area (Figure 4L, p = 0.05), and tended to decay more rapidly (Figure 4M, p = 0.06) in 5xFADs relative to WTs.

**Figure 4.**
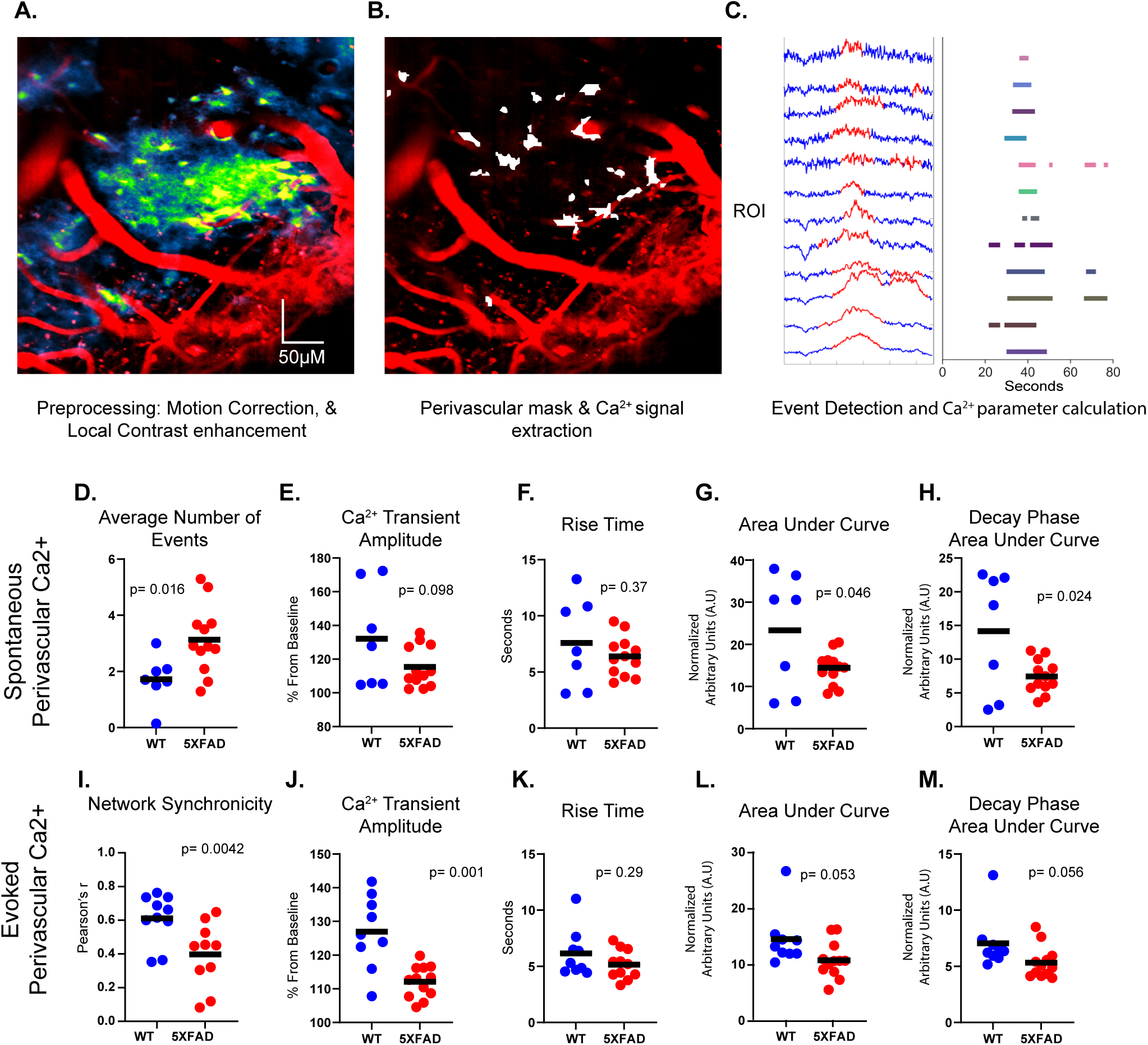
Perivascular astrocyte Ca^2+^ events in awake mice. **A**, Two photon micrograph of cerebral vessels (red) and summated activity map of astrocyte Ca^2+^ signals (green). **B**. Mask of perivascular regions (white) for Ca^2+^ event extraction. **C**. Representative Ca^2+^ traces extracted from perivascular ROIs. In the left panel, regions in red denote where individual Ca^2+^ transients were detected and measured. In the right panel, extracted Ca^2+^ events from unique ROIs were temporally aligned for network analyses, as described in Figure 3B. **D—H**, Scatterplots for spontaneous perivascular Ca^2+^ transient parameters including the average number of transients/ROI (**D**), amplitude (**E**), rise time (**F**), AOC (**G**), and decay (**H**). **I-M**, Scatterplots for air puff-evoked perivascular Ca^2+^ transient parameters including correlated activity across all endfeet pairs in the FOV (**I**, Network synchronicity as described in Figure 3), and average Ca^2+^ transient amplitude (**J**), rise time (**K**), AOC (**L**), and decay (**M**) in individual perviascular ROIs within the FOV. Each plot symbol represents an individual mouse. *p* values derived from two-tailed T tests.

### Reduced signaling fidelity between astrocyte endfeet and arterioles in 5xFAD mice

To investigate dynamic interactions specifically between cerebral arterioles (see Figure 2) and astrocyte endfeet we used a custom MATLAB-based application called LAVA (see Extended data 3 and Figure 5). Analyses were restricted to endfeet pairings with cerebral arterioles because of the relative difficulty of observing/measuring dilations in veins/venules and capillaries. Figure 5A shows a cartoon illustration of arteriole segments demarcated by blue lines along with adjacent astrocyte endfeet. Arrows indicate expansion/constriction of vessel diameter with air puff stimulation. Figure 5B shows a two photon micrograph of an arteriole (red) and summated astrocyte Ca^2+^ activity (green) in the FOV. Arterioles of interest are binarized and linearized (Figure 5C) and immediately adjacent perivascular Ca^2+^ events (1 and 2, shown in green on either side of linearized vessel) are extracted and mapped back to the raw channels (Figure 5D, vessel segment shown in white, adjacent endfeet compartments 1 and 2 shown in fuchsia). Figure E shows the temporal relationships between vasoactive responses observed in the arteriole segment and immediately adjacent astrocyte endfeet shown in Figure 5D. In nearly all arteriole endfeet pairings, Ca^2+^ changes in astrocyte endfeet lagged behind arteriole dilations (also see Extended data 3). A total of 55 and 74 endfeet-arteriole segment pairings were evaluated in the barrel cortex of 10 WT and 13 5xFAD mice, respectively (Figure 5F, 5G). For the WT group, Ca^2+^ transients occurred in 80% of the endfeet in response to a stimulation-induced dilation of the immediately adjacent arteriole segment (Figure 5F). For approximately 18% of the endfeet-arteriole pairings in the WT group, dilations occurred in the absence of Ca^2+^ transients, while in roughly 2% of the pairings there was neither a dilation, nor a Ca^2+^ change (Figure 5F). In contrast to WT mice, an evoked Ca^2+^ transient was observed in only 58% of the endfeet-arteriole pairings in the 5xFAD group (Figure 5G). In 32% of the pairings, arteriole dilations occurred in the absence of an endfoot Ca^2+^ transient, while approximately 4% the pairings showed an endfoot Ca^2+^ response in the absence of an arteriole dilation (Figure 5F). The remaining pairings (5.41%) in the 5xFAD group showed neither a dilation nor a Ca^2+^ change. In 5xFAD pairings where there was co-activity, endfeet Ca^2+^ transients emerged more slowly after local arteriole dilations (Figure 5H, p = 0.027) and were smaller in amplitude (Figure 5I, p = 0.003). We next investigated whether Ca^2+^ changes in astrocyte endfeet were proportional to the magnitude of the arteriole dilation (Figures 5J). For WT mice, endfeet Ca^2+^ transient amplitudes were positively correlated to dilation magnitudes observed in immediately adjacent arterioles (r = 0.38, p = 0.0062). In contrast, we observed no correlation between arteriole dilation magnitudes and endfeet Ca^2+^ changes in 5xFAD mice (r = 0.046, p = 0.71; 5xFAD vs. WT, z = 1.851, p = 0.032). Together, these results suggest that endfeet Ca^2+^ signaling is relatively uncoupled from local arteriole activity in 5xFAD vs. WT mice.

**Figure 5.**
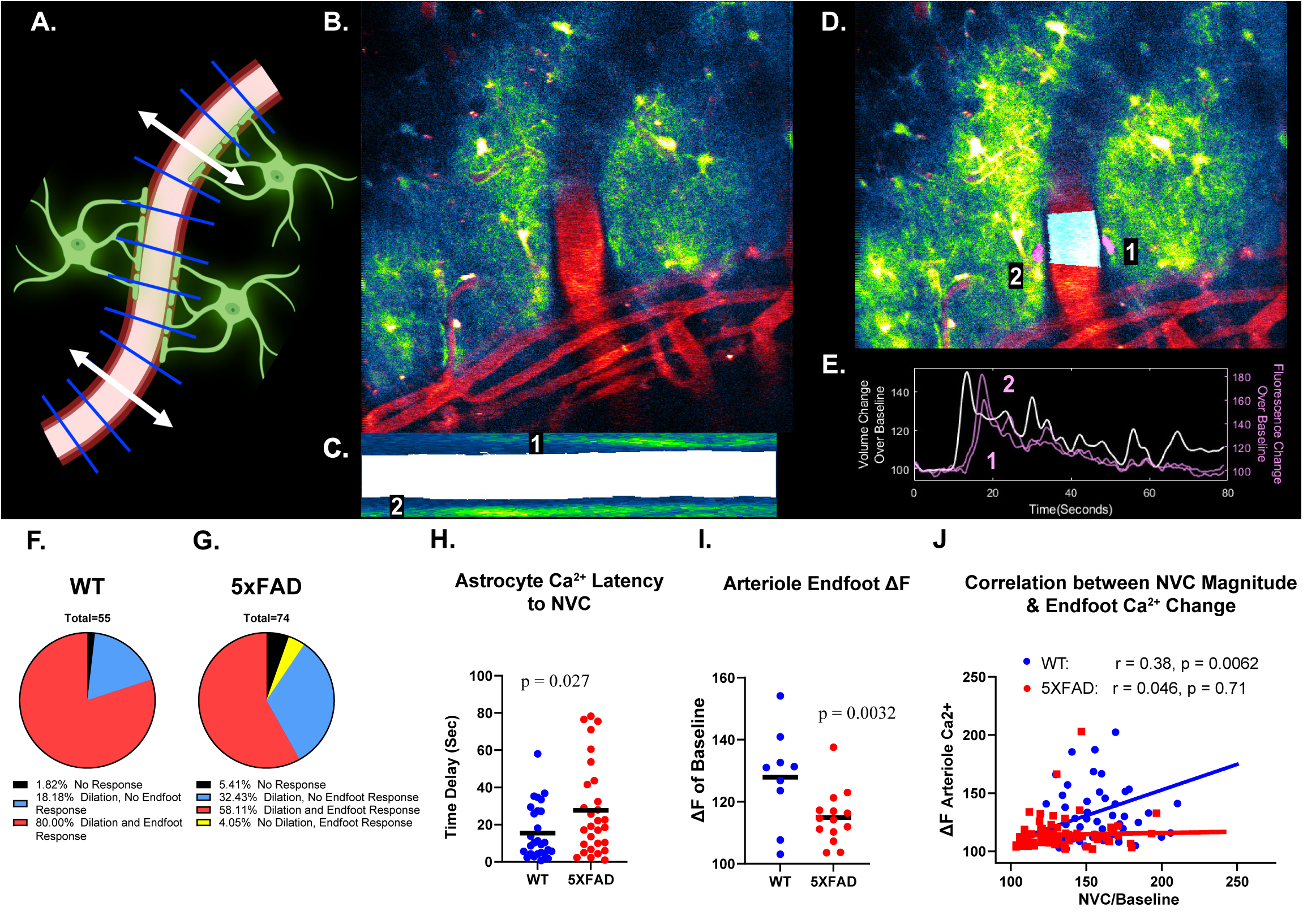
Spatial-temporal analyses of vascular-astrocyte endfeet activity by LAVA application. **A**. Cartoon illustration showing analyses of astrocyte endfeet pairings within individual arteriole segments (blue lines) during airpuff-induced dilations (white arrows). Created in BioRender. Weiss, B. (2025) https://BioRender.com/u91p773 **B**, Two photon micrograph of a cerebral penetrating arteriole (red) and summated activity map of astrocyte Ca^2+^ signals (green). **C**, Modeling and sampling of arteriole crosssections are used to produce linearized channel clips containing the vessel of interest (white) and perivascular spaces (Ca^2+^ activity in green). Vascular tone is measured from cross sectional area measurements, as well as close capture of astrocyte endfeet (1 and 2). **D**, Same FOV shown in panel B assessed during whisker stimulation. Endfoot ROIs (1 and 2 pink) and vascular mask (white) determined from the linearized vessel model (panel C) are mapped onto raw channels. **E**, Measurements of vascular tone from linearlized model in panel C and Ca2+ levels recorded from each endfoot (1 and 2) in panel D are plotted to display temporal associations of field activity. **F-G**, Pie charts showing the functional make-up of endfeet-arteriole pairings for WT (**F**) and 5xFAD (**G**) mice. Red indicates arteriole dilations with a subsequent endfoot Ca^2+^ response; Blue indicates arteriole dilations in the absence of an endfoot response; Yellow indicates endfoot Ca^2+^ response in the absence of an arteriole dilation; Black indicates pairings with neither an arteriole dilation, nor an endfoot Ca^2+^ response. **H**, Scatter plots showing the endfoot Ca^2+^ response latency (*i.e.* latency between initial stimulation induced vasodilation and subsequent endfoot Ca^2+^ signals). **I**, Scatterplot shows evoked Ca^2+^ transient amplitudes in arteriole-associated endfeet. **J**, Scatterplots show the proportional relationship between Ca^2+^ transient amplitude in each endfoot relative to vessel cross sectional area in the immediately adjacent arteriole. In **H** and **J**, each plot symbol represents an individual endfoot-arteriole pairing. In **I**, each plot symbol represents an individual mouse. *p* values derived from two-tailed T tests.

## DISCUSSION

This study is the first we know of that has investigated astrocyte Ca^2+^ signaling in conjunction with arteriole dilatory responses in an intact, and fully awake, mouse model of AD. The overall results demonstrate signs of spontaneous hyperactivity in reactive astrocytes of 5xFAD mice, consistent with other reports. In contrast, evoked Ca^2+^ activity in 5xFAD mice was impaired in individual astrocyte compartments and also across astrocyte networks. Most strikingly, we found that astrocyte endfeet Ca^2+^ signaling in 5xFAD mice is largely uncoupled from local arteriole dilations, which could have important implications for brain metabolism during the progression of AD-related pathology. The results underscore the sensitivity of astrocyte endfeet to developing pathological conditions, like AD, and suggest that these cellular specializations may offer modifiable targets for improving blood flow, metabolism, and overall brain health.

In human brains with abundant AD neuropathology (Aβ plaques and neurofibrillary tangles), there have been reports of comorbid cerebral small vessel pathologies other than cerebral amyloid angiopathy, as well as *in vivo* neuroimaging abnormalities (*e.g.* white matter abnormalities and vascular flow perturbations) indicative of small vessel disease (SVD)(Hilal et al., 2017; Koncz and Sachdev, 2018; Ali et al., 2023; Swinford et al., 2023; Wu et al., 2023; Edwards et al., 2024; Song et al., 2024; Zhang et al., 2024). These observations imply the existence of pathogenetic synergies between AD-type neuropathology and SVD that go well beyond amyloid angiopathy, and which may relate to dysfunction of astrocytes. Ca^2+^ signaling in astrocytes has been linked to numerous processes including the modulation of transmembrane K^+^ channels, release of gliotransmitters, cytoskeletal remodeling/degeneration, and gene expression to name a few(Straub and Nelson, 2007; Sompol and Norris, 2018; Verkhratsky, 2019; Lim et al., 2021). All of these processes appear to be altered in neurodegenerative conditions(Olabarria et al., 2010; Bellot-Saez et al., 2017; Price et al., 2021). Moreover, targeted inhibition of hyperactive Ca^2+^ -signaling pathways in astrocytes ameliorates multiple pathophysiologic outcomes in intact AD/ADRD mouse models(Furman et al., 2012; Sompol et al., 2017; Sompol et al., 2023), suggesting a central role for astrocytic Ca^2+^ abnormalities in chronic neurodegenerative conditions.

To address potential changes in astrocyte Ca^2+^ signaling in the context of AD-like pathology an ever-increasing number of studies has turned to imaging approaches in primary cultures(Lim et al., 2013; Ronco et al., 2014; Mitroshina et al., 2022), *ex vivo* brain slices(Huffels et al., 2022; Paumier et al., 2022; Abghari et al., 2023), and intact animals(Takano et al., 2007; Kuchibhotla et al., 2009; Delekate et al., 2014; Lines et al., 2022; Shah et al., 2022; Kelly et al., 2023; Lee et al., 2023; Sompol et al., 2023). But, while astrocyte-blood vessel interactions are one of the most important, and most direct, ways that the periphery communicates with the brain, few studies have investigated whether Ca^2+^ signaling in reactive astrocytes retains fidelity with cerebral vessel activity. The choice of experimental preparation to address this issue is an important consideration because astrocyte morphologic features and cellular functions depend critically on the presence of adjoining blood vessels and neurons(Lange et al., 2012; Takano et al., 2014). Even in intact animals, the use of anesthesia may alter vascular tone and reduce astrocyte activity, which could uncouple astrocytes from their normal metabolic duties(Thrane et al., 2012; Gao et al., 2017). For these reasons, we chose to investigate astrocyte-blood vessel interactions in fully awake 5xFAD mice at an age where glial reactivity is extensive and neural function is significantly compromised.

### Spontaneous Ca^2+^ activity is augmented in 5xFAD astrocytes, while evoked activity is impaired

Similar to previous reports(Kuchibhotla et al., 2009; Delekate et al., 2014; Takano et al., 2014; Abjorsbraten et al., 2022; Huffels et al., 2022; Lines et al., 2022), 5xFAD mice exhibited an increase in the frequency of spontaneous Ca^2+^ transients across cellular compartments, suggesting that reactive astrocytes are somewhat hyperactive. Increased rates of Ca^2+^ activity may explain why Ca^2+^-dependent pathways, like calcineurin/NFAT, show aberrantly high activity/expression in reactive astrocytes(Lim et al., 2013; Pleiss et al., 2016; Sompol et al., 2017; Kraner et al., 2024). The source of Ca^2+^ hyperactivity may be an intrinsic property of astrocytes, as previous studies observed greater Ca^2+^ transient frequency in APP/PS1 transgenic mice even when neuronal activity was independently suppressed(Kuchibhotla et al., 2009). While astrocytes in 5xFADs appeared spontaneously hyperactive, evoked responses were impaired at the network level and also at the levels of individual cellular compartments. In both transgene groups, whisker stimulation elicited robust astrocytic Ca^2+^ responses that were highly correlated across multiple cells in the field of view. However, correlated activity in 5xFAD mice was significantly lower compared to WTs, perhaps because of gap junction abnormalities that disrupt electrotonic coupling across astrocyte networks(Semyanov, 2019). Reductions in evoked Ca^2+^ transient amplitudes in 5xFAD mice were also observed in astrocyte somata, processes, and endfeet, similar to what others have reported(Lines et al., 2022; Abghari et al., 2023).

### Loss of signaling fidelity between endfeet and arterioles in 5xFAD mice

A highly novel observation of the present work is that astrocyte endfeet in 5xFAD mice responded to penetrating arteriole dilations with poor fidelity. To evaluate concurrent cerebral vessel and astrocyte endfeet activity, we developed a custom MATLAB based application called LAVA. The application is unique in its versatility for analyzing otherwise challenging video files. Its preprocessing options for motion correction, filtering, dissection by trial phases, and adaptive ROI identification allow for precise assessment of vascular tone and proximal Ca^2+^ signals, even for narrow, moving endfeet exhibiting small Ca^2+^ signals. In addition, LAVA enables comparison of its outputs to existing ROI signal analysis. Perhaps the biggest advantage of LAVA is that it can determine the temporal and proportional properties of perivascular Ca^2+^ signals in relation to the activity of immediately adjacent vessel segments. This capability is particularly useful for defining the directional nature of signaling within the neurovascular unit. For instance, under the conditions employed here, we found that evoked Ca^2+^ transients in astrocyte endfeet almost always followed the dilation of local arterioles (Figure 5 and see Extended data 3). These observations are similar to earlier work on fully awake mice during air-puff whisker stimulation, or stimulation from natural whisking during locomotor activity(Tran et al., 2018; Del Franco et al., 2022). While we cannot rule out the possibility that faster events at astrocyte endfeet (*e.g.* transmembrane K^+^ fluxes) contribute to rapid vessel dilations, the LAVA analyses conducted here, in combination with the use of the most sensitive GCaMP available (jGCaMP8f), suggests that astrocyte Ca^2+^ signaling does not initiate local arteriole dilations.

Regardless of whether astrocytes trigger dilation, our work suggests that reactive astrocytes in 5xFAD mice do not respond to arteriole dilations with the same fidelity seen in WT counterparts. Ca^2+^ transients in 5xFAD endfeet occurred only 58% of the time in response to local arteriole dilations (vs 80% in WT mice), with a lag time of about 30 sec (vs. 17 sec in WTs). While 5xFAD mice also exhibited smaller evoked arteriole dilations, it’s unlikely this deficit was the primary cause of impaired endfeet Ca^2+^ signaling. In fact, we found no correlation between the magnitude of vessel dilation and endfeet Ca^2+^ transient amplitude in 5xFAD mice (unlike WTs, which did exhibit proportionality between arteriole and endfeet responses). Together, these results suggest that astrocytes, a major metabolic hub in the brain, respond sluggishly and disproportionately in 5xFAD mice to the most important mechanism for delivering energy on demand (*i.e.* functional hyperemia).

## Limitations and Future Directions

Future work will be necessary to determine why astrocyte Ca^2+^ transients are smaller and respond to vascular activity with lower fidelity in 5xFAD mice. One possibility is a lack of neuronal drive. Like many amyloid models of AD, 5xFAD mice exhibit smaller evoked synaptic responses compared to WT counterparts(Sompol et al., 2017; Forner et al., 2021; Chen et al., 2022), leading to a possible reduction in neuron-astrocyte communication. There is also extensive evidence for structural abnormalities in astrocyte endfeet in AD/ADRD and AD/ADRD animal models. Commonly reported changes include endfeet degeneration, detachment of endfeet from cerebral vessels, and mislocalization of key anchoring proteins like dystrophins(Okoye and Watanabe, 1982; Wilcock et al., 2009; Hawkes et al., 2013; Kimbrough et al., 2015; Sudduth et al., 2017; Abbrescia et al., 2024). Any of these changes could disrupt astrocyte signaling in particular, and brain metabolism in general. It’s also possible that the evoked Ca^2+^ transient deficits we observed here in 5xFAD astrocytes actually reflect a ceiling effect due to elevated resting Ca^2+^ levels. A limitation of our approach is that absolute Ca^2+^ concentrations are difficult to assess with GCaMPs. However, a recent study (also conducted on fully awake mice), using a ratiometric indicator, did report higher resting astrocyte Ca^2+^ levels in a similar amyloid mouse model (Kelly et al., 2023). If basal Ca^2+^ is elevated, this could additionally lead to augmented Ca^2+^ dependent K^+^ fluxes at astrocyte endfeet, which could affect cerebral vessel tone. Finally, cerebral vessels can communicate directly with astrocyte endfeet through Ca^2+^ permeable mechanosensitive channels like TRPV4, which can, in turn, regulate vascular tone directly, or indirectly through vasculo-neuronal coupling(Kim et al., 2015; Haidey et al., 2021). The dysregulation of TRPV4 at astrocyte endfeet could therefore be an additional source for the endfeet/arteriole uncoupling we report here.

## Conclusions

Our results show that reactive astrocytes in a fully awake mouse model of AD-like amyloid pathology are spontaneously hyperactive and exhibit impaired functional connectivity. Changes in Ca2+ signaling observed here are consistent with the Ca2+ hypothesis of aging and AD, which suggests that Ca^2+^ dysregulation in multiple neural types is a driving force in neurodegenerative diseases. (Bezprozvanny and Mattson; Landfield; Sama and Norris; Schrank et al.; Thibault et al.; Lim et al., 2021) Perhaps most notable, we found that astrocyte endfeet respond to dilations in immediately adjacent arterioles with poor fidelity indicating a key point of communication breakdown between the brain and the cerebrovasculature.

## List of abbreviations

AAV, adeno-associated virus; AD, Alzheimer’s disease; ADRDs, AD-related dementias; APP, amyloid precursor protein; AUC, area under the curve; GFAP, glial fibrillary acidic protein; FOV, field of view; LAVA, localized analysis of vascular astrocytes; PS1, presenilin 1; ROI, region of interest; SVD, small vessel disease; TRPV4, ransient receptor potential vanilloid 4; WT wild-type;

## Acknowledgments

Work was supported by National Institutes of Health (NIH)–National Institute on Aging Grants AG078116 to C.M.N., Y.K., Y.J., P.T.N., D.M.W., O.T. and P.S. and AG027297 C.M.N; the Hazel Embry Research Trust; and the Sylvia Mansbach Endowment for Alzheimer’s Disease Research.

